# Roary: Rapid large-scale prokaryote pan genome analysis

**DOI:** 10.1101/019315

**Authors:** Andrew J. Page, Carla A. Cummins, Martin Hunt, Vanessa K. Wong, Sandra Reuter, Matthew T. G. Holden, Maria Fookes, Jacqueline A. Keane, Julian Parkhill

## Abstract

**Summary:** A typical prokaryote population sequencing study can now consist of hundreds or thousands of isolates. Interrogating these datasets can provide detailed insights into the genetic structure of of prokaryotic genomes. We introduce Roary, a tool that rapidly builds large-scale pan genomes, identifying the core and dispensable accessory genes. Roary makes construction of the pan genome of thousands of prokaryote samples possible on a standard desktop without compromising on the accuracy of results. Using a single CPU Roary can produce a pan genome consisting of 1000 isolates in 4.5 hours using 13 GB of RAM, with further speedups possible using multiple processors.

**Availability and implementation:** Roary is implemented in Perl and is freely available under an open source GPLv3 license from http://sanger-pathogens.github.io/Roary

**Supplementary information:** Supplementary data are available at *Bioinformatics* online.

## 1 INTRODUCTION

The term microbial pan genome was first used in 2005 (Medini *et al.*, 2005) to describe the union of genes shared by genomes of interest (Vernikos *et al.*, 2014). Since then, availability of microbial sequencing data has grown exponentially. Aligning whole genome sequenced isolates to a single reference genome can fail to incorporate non-reference sequences. By using *de novo* assemblies, non-reference sequence can also be analysed. Microbial organisms can rapidly acquire genes from other organisms that can increase virulence or promote antimicrobial drug resistance (Medini *et al.*, 2005). Gaining a better picture of the conserved genes of an organism, and the dispensable accessory genome, can lead to a better understanding of key processes such as selection and evolution.

The construction of a pan genome is NP-hard (Nguyen *et al.*, 2014) with additional difficulties from real data due to contamination, fragmented assemblies and poor annotation. Therefore any approach must employ heuristics in order to produce a pan genome (reviewed in (Vernikos *et al.*, 2014)). The most complete standalone pan genome tools are: PanOCT (Fouts *et al.*, 2012), which uses conserved gene neighborhood in addition to homology to accurately place proteins into orthologous clusters; and PGAP which takes annotated assemblies, performs an all-against-all BLAST, clusters the results, and produces a pan genome (Zhao *et al.*, 2012).

PanOCT and PGAP require an all-against-all comparison using BLAST, which means the running time grows non-linearly with the size of input data and is computationally infeasible with large datasets. They also have non-linear memory requirements, quickly exceeding the RAM available in high performance servers for large datasets. We have developed a method to generate the pan genome of a set of related prokaryotic isolates. It works with thousands of isolates in a computationally feasible time, beginning with annotated fragmented *de novo* assemblies. We address the computational issues by performing a rapid clustering of highly similar sequences, which can reduce the running time of BLAST substantially, and carefully manage RAM usage so that it increases linearly, both of which make it possible to analyse datasets with thousands of samples using commonly available computing hardware.

## 2 DESCRIPTION

The input to Roary is one annotated assembly per sample in GFF3 format (Stein, 2013), such as that produced by Prokka (Seemann, 2014), where all samples are from the same species. Coding regions are extracted from the input and converted to protein sequences, filtered to remove partial sequences and iteratively pre-clustered with CD-HIT (Fu *et al.*, 2012). This results in a substantially reduced set of protein sequences. An all-against-all comparison is performed with BLASTP on the reduced sequences with a user defined percentage sequence identity (default 98%). Low complexity regions are masked out (Camacho *et al.*, 2008). Sequences are then clustered with MCL (Enright *et al.*, 2002) and finally the pre-clustering results from CD-HIT are merged together with the results of MCL. Using conserved gene neighbourhood information, homologous groups containing paralogs are split into groups of true orthologs. A graph is constructed of the relationships of the clusters based on the order of occurrence in the input sequences, allowing for the clusters to be ordered and thus providing context for each gene. Full details of the method and outputs are provided in the Supplementary Material.

## 3 RESULTS

We evaluated the accuracy, running time and memory usage of Roary against two similar standalone pan genome applications. In each case we performed the analysis using a single processor (AMD Opteron 6272), and provided 60GB of RAM. Analysis was halted after two days if the applications failed to return a result for the given dataset. We constructed a simulated dataset based on *Salmonella enterica* serovar Typhi (*S. typhi*) CT18, accession AL513382 allowing us to accurately assess the quality of the clustering. We created 12 genomes with 994 identical core genes and 23 accessory genes in varying combinations. All of the applications created clusters that are within 1% of the expected results, with Roary correctly building all genes as shown in Table 1. The overlap of the clusters is virtually identical in all applications.

**Table 1.** Accuracy of each pan genome application on a dataset of simulated data

|  | Core Genes | Total Genes | Incorrect split | Incorrect merge |
| --- | --- | --- | --- | --- |
| **Expected** | **994** | **1017** | **0** | **0** |
| PGAP | 991 | 1012 | 0 | 4 |
| PanOCT | 993 | 1015 | 1 | 1 |
| Roary | 994 | 1017 | 0 | 0 |

In addition, a set of 1000 real annotated assemblies of *S. typhi* genomes was used. Subsets of the data were provided to each application and the running time and memory usage were noted. The running time of PGAP and PanOCT increases substantially, making only small datasets computationally feasible (Fig. 1 and Sup. Fig. 1-8). Roary scales consistently as more samples are added (Sup. Fig. 1-8) and has been shown to work on a dataset of 1000 isolates as shown in Table 2. The memory usage of PGAP and PanOCT also increases rapidly as more samples are added, quickly exceeding 60GB for even small datasets. The memory usage of Roary scales consistently as more samples are added, making it feasible to process large data sets on a standard desktop computer within a few hours. We conducted similar experiments with more diverse datasets including *S. pneumonia*, *S. aureus*, and *Y. enterocolitica* and the results show/exhibit similar speed-ups as shown in Sup Figs 7-8.

**Fig. 1.**
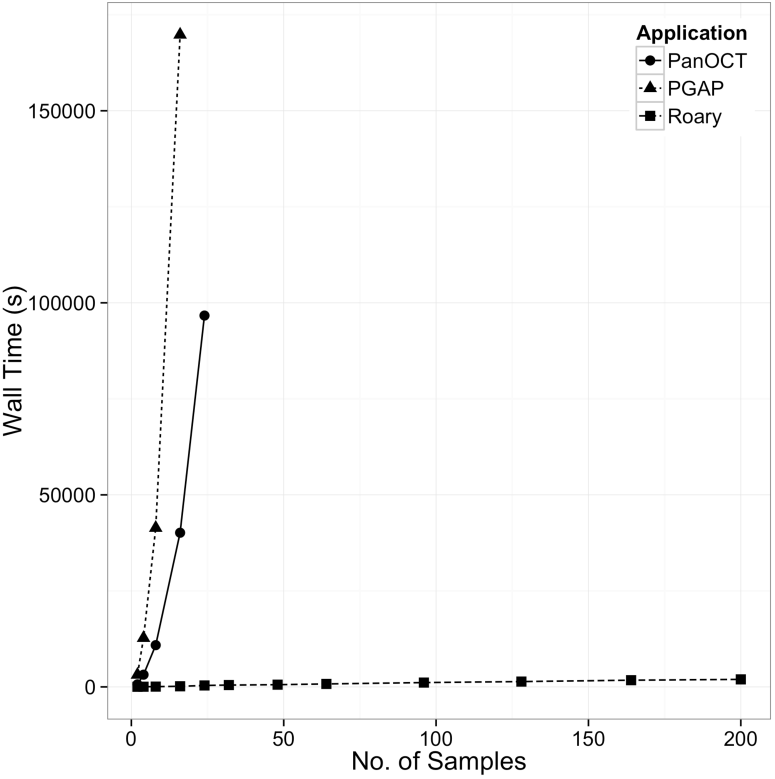
Effect of dataset size on the wall time of multiple applications.

**Table 2.** Comparison of pan genome applications using real *S. typhi* data (ERP001718).

| Samples |  | Core* | Total | RAM (mb) | Wall time (s) |
| --- | --- | --- | --- | --- | --- |
| 8 | PGAP | 4545 | 4929 | 569 | 41397 |
|  | PanOCT | 4544 | 4936 | 663 | 1457 |
|  | Roary | 4459 | 4871 | 179 | 73 |
| 24 | PGAP | - | - | - | - |
|  | PanOCT | 4522 | 4991 | 5313 | 96093 |
|  | Roary | 4436 | 4941 | 349 | 369 |
| 1000 | PGAP | - | - | - | - |
|  | PanOCT | - | - | - | - |
|  | Roary | 4016 | 9201 | 13752 | 15465 |

\* Core is defined as a gene being in at least 99% of samples, which allows for some assembly errors in very large datasets. Where there are no results, the applications failed to complete within 2 days or used more than 60GB of RAM. The first column is the number of unique *Salmonella Typhi* genomes in the input set.

## 4 DISCUSSION

We have shown that Roary can construct the pan genomes of large collections of bacterial genomes using a desktop computer, where it was not previously computationally possible with other methods. Further speedups in running time are possible by providing more processors to Roary. On simulated data Roary is the only application to correctly identify all clusters. This increased accuracy comes from using the context provided by conserved gene neighbourhood information. Roary scales well on large real datasets, identifying large numbers of core genes, even in the presence of a varied open pan genome.

## ACKNOWLEDGEMENTS

This work was supported by the Wellcome Trust (grant number WT 098051).

